# Genotypic variation in resource exchange, use, and production traits in the legume-rhizobia mutualism

**DOI:** 10.1101/2023.08.24.554700

**Authors:** McCall B. Calvert, Maliha Hoque, Corlett W. Wood

**Affiliations:** Department of Biology, University of Pennsylvania, Philadelphia, PA, USA

**Keywords:** mutualism evolution, genotype by genotype interaction, resource exchange, resource use, resource production, legume, rhizobia, *Medicago truncatula*, *Enisfer*

## Abstract

Mutualisms, reciprocally beneficial interaction between two or more species, are ubiquitous in nature. A common feature of mutualisms is extensive context-dependent variation in fitness outcomes. This context-dependency is hypothesized to stem from the environment’s mediation of the relative costs and benefits associated with mutualisms. However, traits related to the exchange of goods and services in mutualisms have received little attention in comparison to net fitness outcomes. In this study, we quantified the contribution of host and symbiont genotypes to variation in resource exchange, use, and production traits measured in the host using the model mutualism between legumes and nitrogen-fixing rhizobia. We predicted that plant genotype x rhizobia genotype (GxG) effects would be common to resource exchange traits because resource exchange is hypothesized to be governed by both interacting partners through bargaining. On the other hand, we predicted that plant genotype effects would dominate host resource use and production traits because these traits are only indirectly related to the exchange of resources. Consistent with our prediction for resource exchange traits, but not our prediction for resource use and production traits, we found that rhizobia genotype and GxG effects were the most common sources of variation in the traits that we measured. The results of this study complement the commonly observed phenomenon of GxG effects for fitness by showing that numerous mutualism traits also exhibit GxG variation. Furthermore, our results highlight the possibility that the exchange of resources as well as how partners use and produce traded resources can influence the evolution of mutualistic interactions. Our study lays the groundwork for future work to explore the relationship between resource exchange, use and production traits and fitness (i.e. selection) to test the competing hypotheses proposed to explain the maintenance of fitness variation in mutualisms.

## Introduction

Almost all multicellular life engages in some form of mutualism. Mutualisms are interspecific interactions in which partners increase their fitness by reciprocally exchanging goods and services. Genetic variation for fitness outcomes of mutualisms are notoriously context dependent, presumably because the environment influences the costs and benefits of the interaction for both partners. However, variation in the benefit and cost traits that underlie mutualisms has received comparatively little attention.

Fitness-related traits have been useful for inferring net interaction outcomes. However, the measurement of fitness-related traits also obscures the individual traits that underlie fitness variation. The fitness outcome of a mutualistic interaction is determined by the balance between its benefits and costs. Benefits can take a variety of forms, including transportation (e.g. the movement of gametes in plant-pollinator mutualisms; Albrecht et al. 2012), protection (e.g. defense against herbivory in ant-acacia mutualisms; Janzen 1966), and nutrition (e.g. the exchange of limiting resources between plants and microbes; Friesen et al. 2011). Costs are related to the energetic investment involved in producing benefits that are transferred to the interacting partner (Bronstein 2001, 2015). For example, in plant-microbe mutualisms that involve an exchange of resources, a principal cost for the host is the carbon that is transferred to microbial partners (Akçay and Roughgarden 2007; Clark et al. 2019).

Beyond the traits involved in the direct exchange of resources, traits related to the use or production of the traded resources can also influence the net fitness consequences of a mutualism by impacting its relative benefits and costs (Vidal and Segraves 2021). Traits like resource use efficiency dictate how efficiently the benefits received from an interacting partner are incorporated into physiological processes (e.g. growth, reproduction, defense) that are linked to fitness. Resource production traits, on the other hand, are involved in the creation of the goods and services that are traded to an interacting partner. These traits dictate the total resource pool available to invest in mutualism, and thereby affect the proportional cost of resource exchange. In plant-microbe interactions, a common resource production trait is photosynthetic rate, because it determines the size of a plant’s carbon pool, a portion of which is traded to microbial partners (Grman et al. 2012; Clark et al. 2019).

Like other phenotypes, variation in resource exchange, use, and production traits is likely attributable to genetic variation in one or both partners. Indeed, partner genotype effects, where the fitness outcome of a mutualism depends on the genotype of one or both of the interacting partners, are a common feature of resource mutualisms (Hoeksema and Bruna 2015). An added layer of complication is that traits mediating species interactions can be jointly determined by both interacting partners (Moore et al. 1997; O’Brien et al. 2021b; De Lisle et al. 2022). Traits are jointly determined when variation in a trait is determined by contributions from the genomes of both interacting partners. Therefore, genetic variation in traits that mediate a host-microbe mutualism could arise from host genotype effects, microbe genotype effects, or from genotype-by-genotype (GxG) interactions (Heath 2010; Friesen et al. 2011).

Despite their ecological and evolutionary significance, however, few empirical studies have investigated whether there is heritable variation in resource exchange, use, and production traits in interacting populations. Quantifying genotypic differences in mutualism traits represents an important synthesis between the subfields of mutualism physiology, functional ecology, and evolution. The underlying metabolic and genetic pathways involved in mutualism function are well defined in many systems, especially in nutritional mutualisms between hosts and symbionts (Heath et al. 2012; Udvardi and Poole 2013; Levy et al. 2021). Furthermore, there is an extensive body of theoretical work on the resource economics of mutualisms (Akçay and Roughgarden 2007; Akçay and Simms 2011; Werner et al. 2014). However, little work has been done that leverages measurements of traded resources to quantify variation in important mutualism components like resource exchange, use, and production.

In this study, we quantified the contributions of host and symbiont genotype to resource exchange and host resource use and production traits in the model mutualism between legumes and nitrogen-fixing bacteria (rhizobia). The legume-rhizobia mutualism is centered on the exchange of carbon for nitrogen, whereby the legume host trades a portion of its carbon budget to rhizobia in exchange for fixed atmospheric nitrogen (Oldroyd et al. 2011; Udvardi and Poole 2013). Quantitative genetic experiments have established that fitness outcomes in the legume-rhizobia mutualism are highly context-dependent, varying across plant and rhizobia genotypes as well as environmental factors like resource availability (Heath and Tiffin 2007; Heath 2010; Weese et al. 2015; Batstone et al. 2017; Burghardt et al. 2017). However, little is known about how resource exchange, use, and production traits vary in natural populations of legumes and rhizobia.

Here, we performed a factorial experiment in which we inoculated two plant genotypes with two rhizobia genotypes and measured the contributions of plant genotype, rhizobia genotype, and their interaction (GxG) to resource exchange, use, and production traits.

Exchange traits are those directly related to the transfer of nitrogen (benefits) to host plant tissue and the transfer of carbon (costs) to rhizobia. Resource use and production traits are those involved in how host plants allocate the nitrogen received from rhizobia and how efficiently they fix (via photosynthesis) the carbon they transfer to rhizobia. We measured many of these traits using elemental analysis, and as such, these traits represent the cumulative endpoint of the dynamic processes. Given that the outcomes of resource exchange are hypothesized to be governed by bargaining between both interacting partners (Sachs et al. 2004; Akçay and Roughgarden 2007; Clark et al. 2019), we predicted that GxG effects would be common to exchange traits. On the other hand, we predicted that only plant genotype effects would be found for resource use and production traits because those traits represent components of plant physiology that should only be indirectly related to the exchange of resources among partners. We found extensive differences between plant and rhizobia genotypes, as well as GxG interactions, for all of the plant traits we measured. Our results suggest that multiple resource-related traits likely contribute to variation in mutualism fitness outcomes.

## Methods

### Study system

The mutualism between *Medicago truncatula* and *Ensifer* spp. is a deeply-studied model system that has been used to uncover the molecular and genetic basis of symbiotic interactions and to explore the evolution of mutualisms (Udvardi and Poole 2013). *M. truncatula* is an annual plant native to the Mediterranean Basin (Urban 1873; Lesins and Lesins 1979). *M. truncatula* forms an endosymbiotic interaction with rhizobia species belonging to the genus *Ensifer* (Andrews and Andrews 2017). This interaction is localized to specialized structures on the host plant’s roots called nodules. As with other legume-rhizobia interactions, *M. truncatula* trades photosynthetically-fixed carbon to rhizobia in nodules in exchange for fixed nitrogen. Previous studies involving the *M. truncatula* - *Ensifer* mutualism have shown that fitness outcomes in this interaction depend on the specific combination of plant and rhizobia genotypes (G x G) (Heath 2010). Past studies using *M. truncatula* have primarily used aboveground biomass as a proxy for fitness.

For this study, we used two inbred lines of *M. truncatula*, HM001 and HM028, sourced from the Medicago HapMap Project (Stanton-Geddes et al. 2013). These accessions are derived from collections made in the natural range of *M. truncatula* in the Mediterranean Basin. We chose these accessions because they, along with many other accessions, display genetic variation for fitness proxies like aboveground biomass (Stanton-Geddes et al. 2013; Wood et al. 2018). We also used two strains of *Ensifer meliloti*, MAG 154 and MAG 282, sourced from a panel of 191 strains isolated from *M. truncatula* sampled from the Mediterranean Basin (Katy Heath, U. Illinois). We selected these strains because they had divergent effects on plant performance when inoculated on *M. truncatula* genotype HM101. As with the inbred lines of *M. truncatula*, we chose these strains because they exhibited variable effects on plant fitness proxies like aboveground biomass (Batstone et al. 2022).

### Trait measurement

We measured a series of resource exchange, use, and production traits on host plants (Table 1). We also measured two proxies for mutualism outcome: plant fitness and nodulation (Table 1). Plant fitness was estimated by weighing the total dry aboveground biomass for an individual plant. We measured three nodule traits: total nodule number, total nodule mass, and average nodule size (total nodule number / total nodule mass). By harvesting plants and measuring aboveground biomass at two different timepoints, we were also able to infer genotype effects on growth rate. *M. truncatula* populations are thought to be subject to viability selection in their dry, Mediterranean habitat, thus growth rate could be an important fitness component (Urban 1873; Lesins and Lesins 1979).

**Table 1:** The mutualism outcome and resource exchange, use, and production traits measured in this study. Brief descriptions of how each trait was and calculated are also provided.

| Mutualism component |  | Trait | Measurement / Calculation |
| --- | --- | --- | --- |
| Outcome | Plant fitness | Aboveground biomass | Dry weight of aboveground plant tissue |
|  |  | Aboveground biomass growth rate | Dry weight of aboveground plant tissue measured at two timepoints (see methods) |
|  | Nodulation | Total nodule mass | Dry weight of all nodules from an individual plant's root system |
|  |  | Nodule number | Count of all nodules on individual plant's root system |
|  |  | Average nodule mass | Total nodule mass / nodule number |
| Exchange traits | Benefit | Nitrogen composition | Proportion of aboveground plant tissue that is composed of nitrogen (measured via elemental analysis) |
|  |  | Total nitrogen benefit | Nitrogen composition of aboveground tissue * aboveground biomass |
|  |  | Total nitrogen benefit per symbiont | Slope of the relationship between total nitrogen benefit and total nodule mass (see "statistical analysis" in methods) |
|  | Cost | Total carbon cost | Carbon composition of nodule tissue * total nodule mass |
|  |  | Nodule carbon fraction | Total carbon cost / (total carbon cost + (aboveground tissue carbon composition * aboveground biomass)) |
|  | Resource exchange ratio | Carbon-for-nitrogen exchange ratio | Total carbon cost / total nitrogen benefit |
| Resource Use | Nitrogen use efficiency | Carbon:nitrogen ratio in aboveground tissue | Nitrogen composition in aboveground plant tissue / carbon composition in aboveground plant tissue |
| Resource Production |  | Leaf photosynthetic rate | See Methods and Supplementary Methods |

Unless otherwise noted, all resource exchange, use, and production traits were measured using carbon and nitrogen elemental composition data collected via combustion analysis in the Department of Earth and Environmental Sciences at the University of Pennsylvania. To perform combustion analysis, tissue was pulverized using either a mechanical ball grinder or a mortar and pestle. Then, approximately 4 mg of pulverized tissue was weighed using an ultra-micro balance (CP2P Microbalance, Sartorius GmbH, Goettingen, Germany), packaged into tin capsules, and ran on a Costech ECS4010 Elemental Analyzer (Costech Analytical Technologies, Inc. Valencia, CA, USA). The elemental analyzer returns measurements of carbon and nitrogen composition (%) for the tissue sample analyzed. We multiplied carbon and nitrogen composition by the total dry mass that the elemental analysis sample came from to infer the amount of total biomass carbon and nitrogen in a given sample (Table 1). We performed combustion analysis separately on one aboveground plant tissue sample and one nodule tissue from between 6 and 10 individual plants from each genotype combination. For elemental analysis on aboveground tissue, we pulverized the entirety of the aboveground portion of the plant, while for elemental analysis on nodules, we pulverized all of the nodules that were dissected from the plants root system.

We measured three benefit-associated exchange traits: nitrogen composition (%) in aboveground tissue (hereafter referred to as nitrogen composition), total nitrogen in aboveground tissue (hereafter referred to as total nitrogen benefit), and total nitrogen in aboveground tissue as a function of total nodule mass (hereafter referred to as total nitrogen benefit per symbiont; see statistical analysis section for more details; Table 1). We measured two cost-associated exchange traits: total carbon in nodule tissue (hereafter referred to as total carbon cost), and the fraction of total carbon in aboveground tissue and nodule tissue that is in nodule tissue (hereafter referred to as nodule carbon fraction). These cost traits incorporate both the carbon cost of constructing nodules as well as the carbon traded to the symbiont. This approach is unable to quantify the portion of carbon transferred to rhizobia that was respired by rhizobia, thus our analysis is making the simplifying assumption that respiration rates are consistent across genotype combinations. We also calculated the ratio of total nitrogen in aboveground tissue to total carbon in nodule tissue as a proxy of the carbon-for-nitrogen exchange ratio.

We measured one resource use trait and one resource production trait. The resource use trait was measured as the ratio of total carbon to total nitrogen in aboveground tissue as a proxy for nitrogen use efficiency (Dovrat et al. 2020). The benefit production trait we measured was leaf photosynthesis rate. Photosynthesis rate measurements were made using a LI-6400 Portable Photosynthesis System (LI-COR, Inc., Lincoln, NE, USA) with an 6400-02B LED Light Source (LI-COR, Inc., Lincoln, NE, USA) attached to the leaf chamber to apply a consistent level of light for all measurements. See the supplementary methods for details on how photosynthesis rate was measured and calculated (Supplementary methods). Table 1 provides short summaries of how each resource exchange, use, and production trait was quantified.

It should be emphasized that resource exchange, use, and production are dynamic and interrelated processes. For example, how efficiently resources are produced can have knock on effects on resource exchange and vice versa. In this study we have designated measured traits as belonging to one of resource exchange, use, and production, but this does not necessarily mean that variation in these traits is related to that process alone. Precise attribution of traits to specific mutualism processes is a general challenge for the field of mutualism evolutionary ecology as a whole and will emerge through more manipulative experiments.

### Experimental design

To quantify the effect of plant genotype, rhizobia genotype, and GxG interactions on the traits we measured, we grew our two *M. truncatula* accessions in fully factorial design with our two *E. meliloti* strains for a total of four plant-rhizobia genotype combinations. Each of the four genotype combinations was replicated 33 times totalling 132 experimental units.

Seeds were scarified, sterilized, imbibed, and stratified using the methods reported in the “Seed storage and germination” chapter of the Medicago Handbook (Noble Institute; Supplementary methods). Prior to planting, we filled 66 mL Cone-tainer pots (Stuewe & Sons, Inc., Tangent, OR, USA) with a growing media consisting of a 1:4 mix of perlite and sand. We sterilized the filled Cone-tainer by autoclaving them twice. One seedling was placed in each Cone-tainer. One day after planting, seedlings were given 1 mL of 0.625 mM nitrogen Fahraeus fertilizer (Batstone et al. 2017). Afterwards, plants were given 0.0 mM N Fahraeus fertilizer every 2 to 3 days to ensure that rhizobia were the only source of nitrogen (Batstone et al. 2017).

Cone-tainer pots were placed in 98-well racks. To avoid the risk of cross contamination of rhizobia strains, every plant in a rack received the same rhizobia strain. Each rack was enclosed in a semi-permeable tent consisting of plastic wrap and PVC pipe to further reduce the risk of cross contamination. Plants were grown in two growth chambers in a complete block design. The height of light fixtures in both chambers was adjusted so that a photosynthetic active radiation (PAR) reading of ∼250 μmol/m^-2^/s^-1^ was consistently measured on the shelf of the growth chambers. Chambers were set to 22°C and a 16:8 light:dark photoperiod. Seedings were maintained in a high humidity environment for about one week, after which they were exposed to ambient humidity, usually 40-50%.

Two weeks after planting, we inoculated plants with rhizobia. Plants received either strain MAG 154 or MAG 282. We grew strains from glycerol stocks in sterile tryptone yeast liquid media in a shaking incubator at 30 °C and 180 rpm for approximately two days. We then streaked the media on sterile Petri plates containing tryptone yeast agar under a laminar flow hood. We incubated the plates at 30°C until colonies started to form. Next, we selected an individual colony from each strain to grow in tryptone yeast liquid media. Again, rhizobia cultures were allowed to grow for approximately two days. We diluted the liquid media to an OD_600_ of 0.1 and inoculated each plant with 1 ml of the diluted culture.

Plants were harvested in three separate cohorts at 9, 10, and 11 weeks post planting. Approximately 20 plants from each treatment group were harvested at 11 weeks. Of those plants harvested at 11 weeks, approximately 10 from each treatment were used to measure the resource exchange, use, and production traits described above via elemental analysis.

Approximately 3 plants from each treatment group were harvested at 10 weeks to measure photosynthesis. Prior to harvest at the 10 week timepoint, photosynthesis rate measurements were taken on 2-3 leaves of each plant. Approximately 10 plants from each treatment group were harvested at 9 weeks. Plants in the 9-week cohort were compared to those in the 11-week cohort to estimate growth rate (see statistical analysis section for more details).

At each harvest, we gently removed plants from their Cone-tainer pots and cut the shoots from the roots. The shoots were placed in brown paper bags and dried at 60 °C for at least a week before being weighed with an analytical scale to measure aboveground biomass. Plants used for benefit and cost trait measurement via elemental analysis were processed as described in the “Trait measurement” section. The roots were washed before being placed in plastic bags and stored in a freezer at -20 °C.

To collect nodulation data, frozen roots were thawed and nodules were removed under a dissecting microscope. Nodules were counted as they were removed to quantify nodule number. To assess the consistency of nodule count estimates across researchers, we had each researcher count nodules on the same subset of 10 plants. Nodule counts were highly repeatable across researchers in a linear mixed effects model with nodule number as the response, and plant and researcher as random effects (researcher explained only 0.4% of variation in nodule counts). Removed nodules were placed in coin envelopes, dried at 60 °C for at least a week, and weighed. Nodules from plants that were used for benefit and cost trait measurement via elemental analysis were processed as described in the “Trait measurement” section.

### Statistical analysis

We used linear models to test for an effect of plant genotype, rhizobia genotype, and plant genotype by rhizobia genotype (GxG) interactions on the traits we measured. All models used a normal error distribution except for the model with nodule number as the response variable, which used a negative binomial error distribution. Block was included as a fixed effect in all models. Continuous predictors were mean-centered to avoid spurious inference on main effects in the presence of interactions (Schielzeth 2010). For some models, response variables were log-transformed to better conform with the assumption of normality. If a response variable was log transformed, it will be indicated in the figure caption of the associated figure. For analysis of elemental composition measurements (e.g. nitrogen composition), arcsine square root transformations were performed to normalize the data prior to running the models. Models with nodule traits as the response included total root mass as a covariate to account for differences in nodule traits associated with differences in root system size.

To test for genotype effects on growth rate, we included the following interaction terms in the model, plant genotype x timepoint, rhizobia genotype x timepoint, and plant genotype x rhizobia genotype x timepoint.

To test for significant genotype effects on total nitrogen benefit per symbiont, we included the interaction terms plant genotype x total nodule mass, rhizobia genotype x total nodule mass, and plant genotype x rhizobia genotype x total nodule mass terms in our models. Each of these interaction terms estimates differences in the slope of the relationship between total nitrogen in aboveground tissue and nodule mass that are due to genotype effects.

Our model with photosynthesis rate as the response included each individual leaf measurement as an independent replicate. To account for any variation due to measurements made on leaves from the same plant, we included plant ID as a random effect.

The significance of model terms in each model was tested using type III sum of squares. For linear models, significant testing was performed using *F*-statistics, and for generalized linear models, significance testing was performed with Wald X^2^ tests.

Many of the traits that we measured are correlated with one another (Fig. S1) These correlations could stem from a physiological relationship or a mathematical relationship resulting from an overlap in the measurements used to calculate a given trait. We caution against drawing robust interpretations from these correlations because of the widespread mathematical non-independence, but we do suggest areas of future research that these correlations could inform (see discussion).

All statistics and data visualization were performed in R v4.2.3 (R Core Team 2023). For simple linear models, we used the base function lm; for linear mixed effects models we used the lmer function in the lme4 package (Bates et al. 2014), and for generalized linear mixed effects models we used the glmmtmb function in the glmmTMB package (Brooks et al. 2017). Model assumptions were evaluated using the DHARMa package (Hartig and Lohse 2022). Significant testing was performed with the Anova function in the car package (Fox and Weisberg 2018). All plots include estimated marginal means that were calculated using the ggeffects package (Lüdecke 2018). The ggplot2 (Wickham 2016), ghibli (Henderson et al. 2022), and ggpubr (Kassambara 2023) packages were used to make all of the plots presented here. All data and code used in this study can be found here [Dryad repository will be added if/when manuscript is accepted for publication]

## Results

### Genetic variation in fitness and mutualism outcomes

We found a significant plant genotype x rhizobia genotype (GxG) interaction effect for aboveground biomass, our proxy for plant fitness, but no main effects of either plant or rhizobia genotype (Fig. 1, Table 2). We also found significant plant genotype, rhizobia genotype, and GxG effects on aboveground biomass growth rate (Fig. 1, Table 2). During the analysis of growth rate, we identified an outlier in the HM028 - MAG 282 - Timepoint 1 treatment group. Removal of this outlier does not change the statistical significance of any model terms.

**Table 2:**
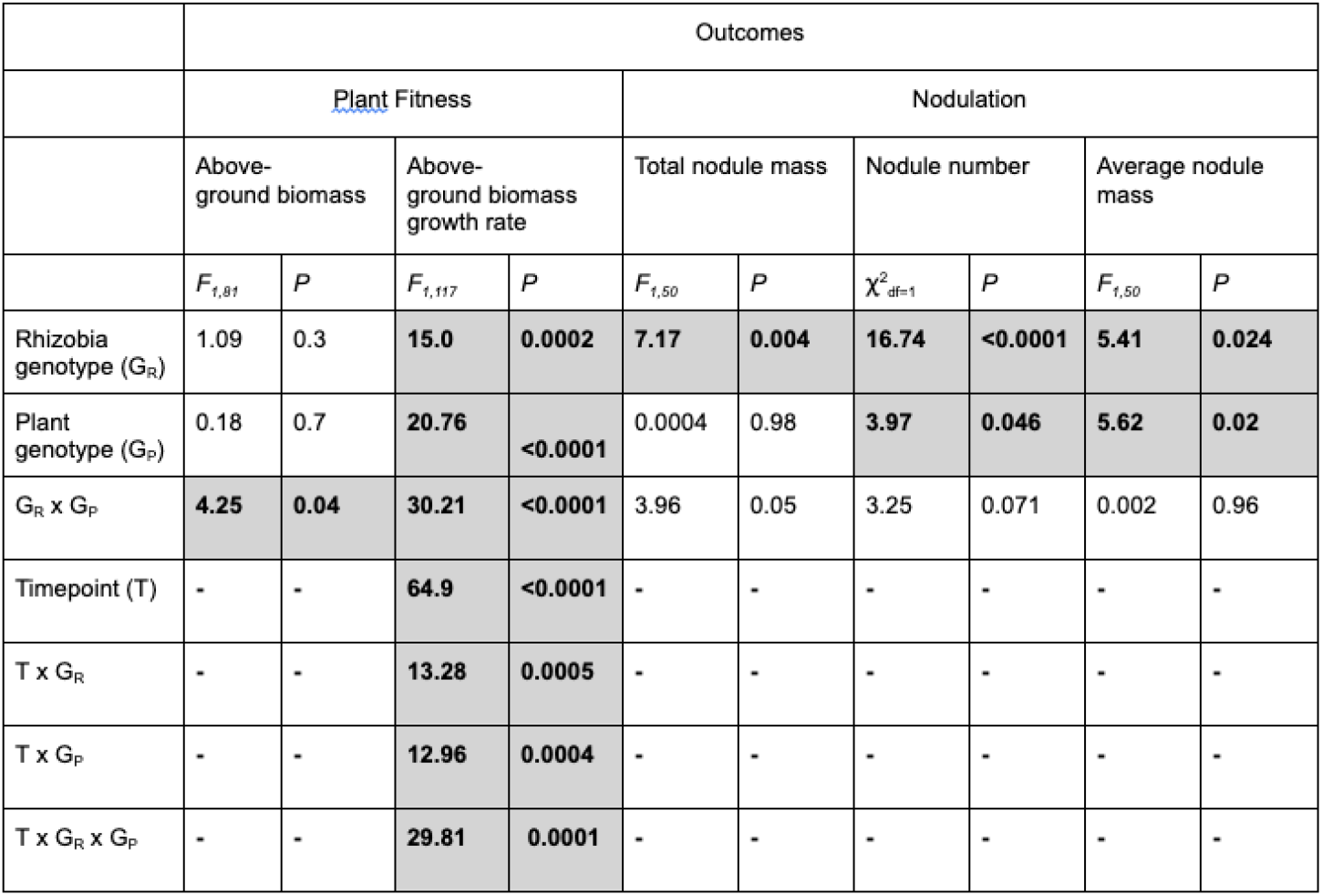
Genotype effects on mutualism outcome traits. Significant model terms are highlighted in gray.

|  | Outcomes |  |  |  |  |  |  |  |  |  |
| --- | --- | --- | --- | --- | --- | --- | --- | --- | --- | --- |
|  | Plant Fitness |  |  |  | Nodulation |  |  |  |  |  |
|  | Above-ground biomass |  | Above-ground biomass growth rate |  | Total nodule mass |  | Nodule number |  | Average nodule mass |  |
| | $F_{1,81}$ | $P$ | $F_{1,117}$ | $P$ | $F_{1,50}$ | $P$ | $\chi^2_{df=1}$ | $P$ | $F_{1,50}$ | $P$ |
| Rhizobia genotype ($G_R$) | 1.09 | 0.3 | 15.0 | 0.0002 | 7.17 | 0.004 | 16.74 | <0.0001 | 5.41 | 0.024 |
| Plant genotype ($G_P$) | 0.18 | 0.7 | 20.76 | <0.0001 | 0.0004 | 0.98 | 3.97 | 0.046 | 5.62 | 0.02 |
| $G_R \times G_P$ | 4.25 | 0.04 | 30.21 | <0.0001 | 3.96 | 0.05 | 3.25 | 0.071 | 0.002 | 0.96 |
| Timepoint (T) | - | - | 64.9 | <0.0001 | - | - | - | - | - | - |
| T x $G_R$ | - | - | 13.28 | 0.0005 | - | - | - | - | - | - |
| T x $G_P$ | - | - | 12.96 | 0.0004 | - | - | - | - | - | - |
| T x $G_R \times G_P$ | - | - | 29.81 | 0.0001 | - | - | - | - | - | - |

**Figure 1:**
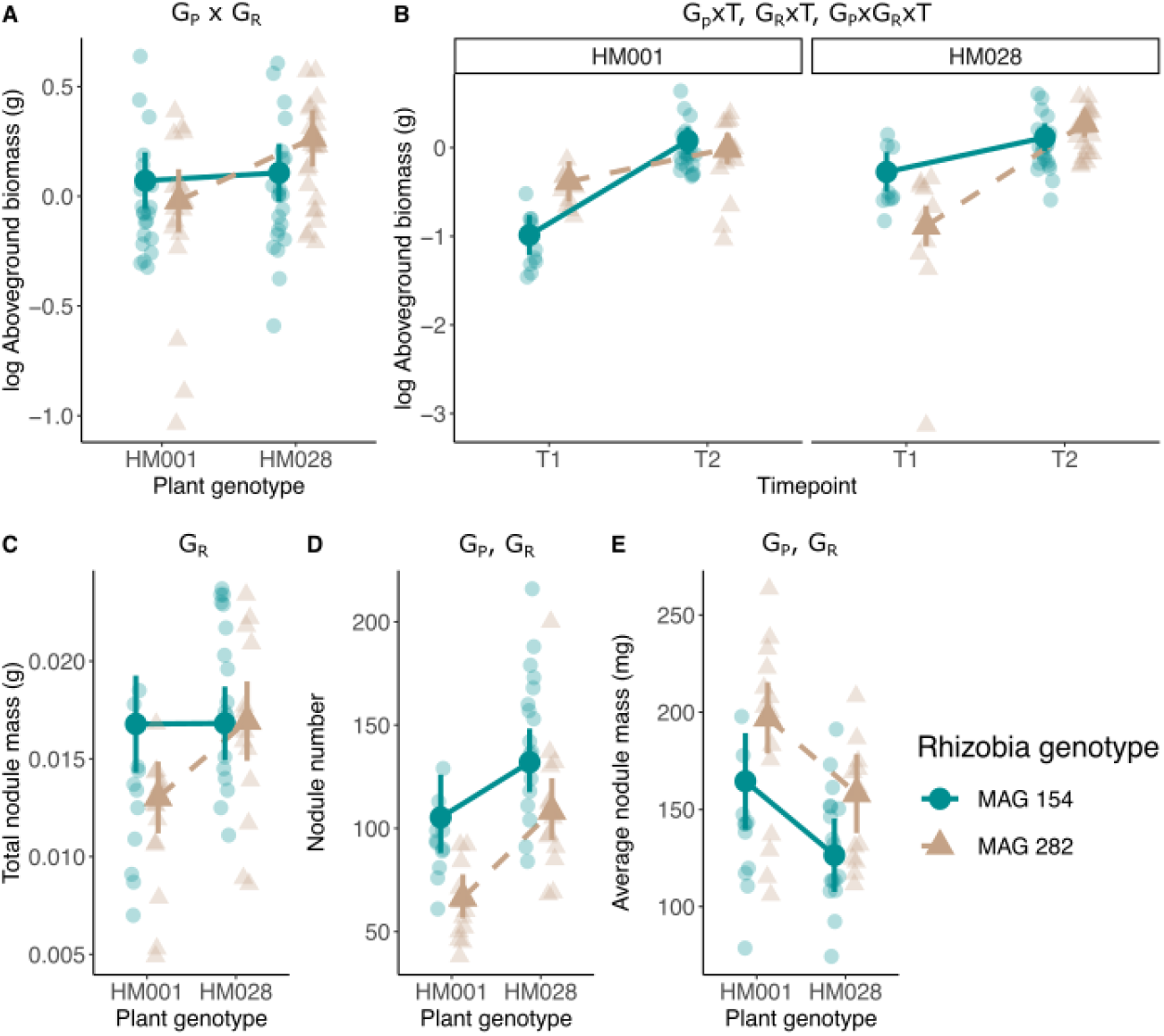
Genetic variation for (A) log transformed aboveground biomass, our proxy for plant fitness, (B) log transformed aboveground biomass growth rate, (C) total nodule mass, (D) nodule number, and **(E)** average nodule size. In all panels, transparent points are the raw data and opaque points are the estimated marginal means. Error bars are the 95% confidence interval. Significant genotype effects are listed on the figure: G_p_= plant genotype effect, G_R_=rhizobia genotype effect, G_p_,x G_R_= plant genotype x rhizobia genotype interaction, G_p_x **T** = plant genotype x timepoint interaction, G_R_x T = rhizobia genotype x timepoint interaction, and G_p_xG_R_x T = plant genotype x rhizobia genotype x timepoint interaction. Model term statistical significance did not change when the outlier in the HM028-MAG282-Timepoint 1 treatment group in panel B was removed.

We observed a significant rhizobia genotype effect on total nodule mass, and significant plant genotype effects and rhizobia genotype effects on total nodule number and average nodule size (Fig. 1, Table 2).

### Genetic variation in exchange traits

We measured three benefit-associated exchange traits, nitrogen composition, total nitrogen benefit, and total nitrogen benefit per symbiont (Table 1). We observed a significant rhizobia genotype effect and GxG effect for nitrogen composition in aboveground tissue (Fig. 2, Table 3), but no significant genotype effects for total nitrogen biomass in aboveground tissue (Fig. 2, Table 3). For total nitrogen per symbiont, we observed significant plant genotype x total nodule mass, rhizobia genotype x total nodule mass, and G x G x total nodule mass effects, indicating that our measure of benefit per symbiont varies in response to plant genotype, rhizobia genotype, and the interaction of plant genotype and rhizobia genotype (Fig. 2, Table 3).

**Table 3:**
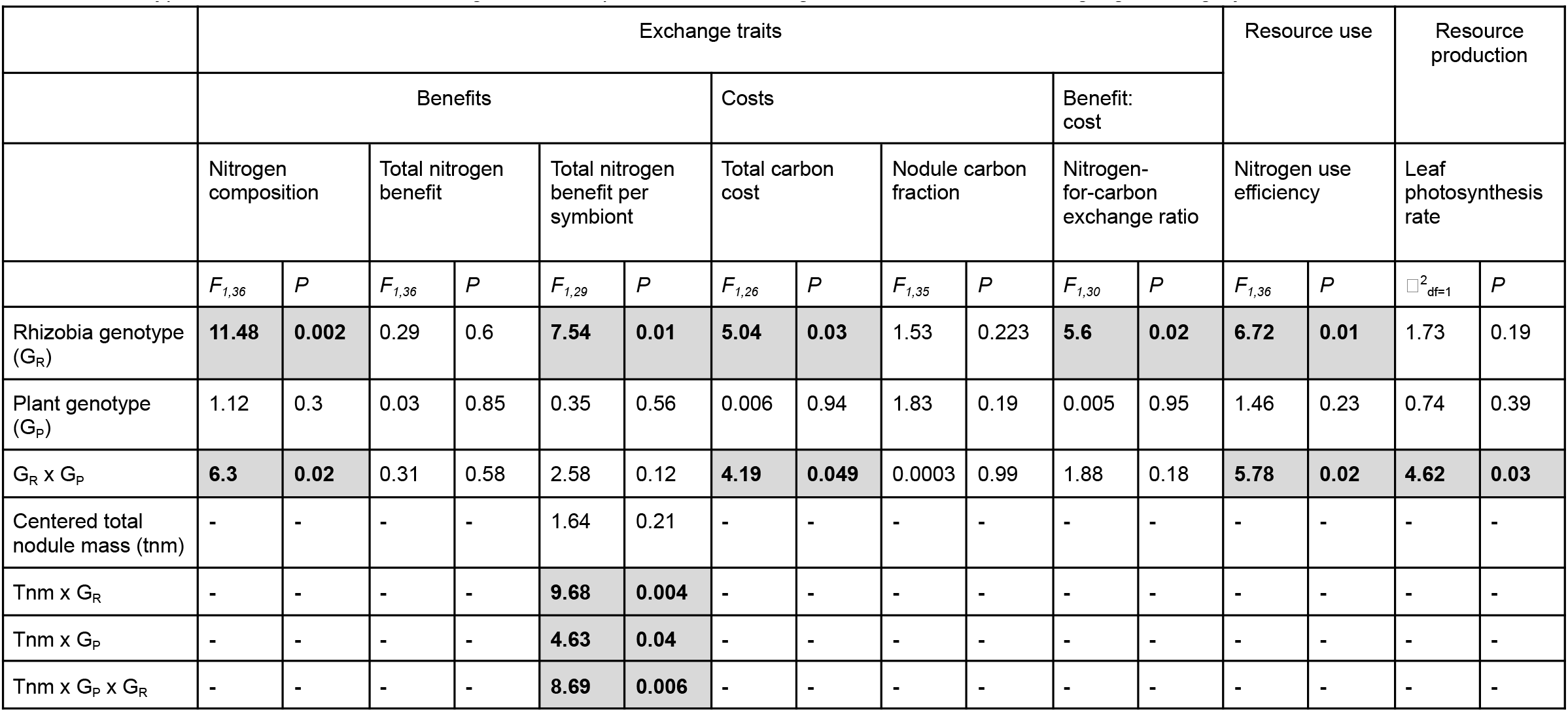
Genotype effects on resource exchange, use, and production traits. Significant model terms are highlighted in gray.

**Figure 2:**
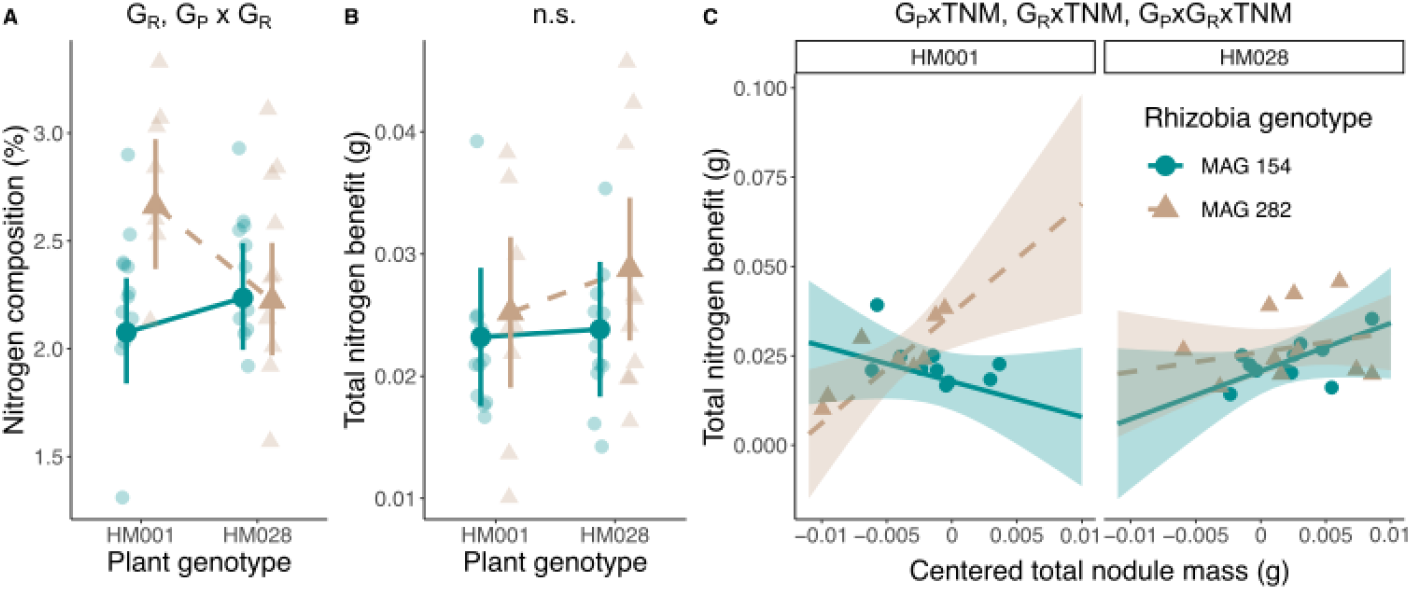
Genetic variation for three benefit-associated exchange traits: (A) nitrogen composition (%), (B) total nitrogen benefit (g), and (C) total nitrogen benefit per symbiont. In panels A and B, transparent points are the raw data, opaque points are the estimated marginal means, and Error bars are 95% confidence intervals. For panel C. transparent points are the raw data, lines represent the estimated marginal slopes derived from the model, and ribbons are the 95% confidence interval for the slope. Significant genotype effects are listed on the figure: G_P_ = planet genotype effect, G_R_ = rhizobia genotype effect, G_P_xG_R_ = plant genotype x rhizobia genotype interaction, G_P_xTNM, G_R_xTNM, G_P_xG_R_xTNM.

We measured two cost-associated exchange traits, total carbon cost, and the nodule carbon fraction (Table 1). We observed a significant rhizobia genotype and GxG effect for total carbon cost, but no significant genotype effects for nodule carbon fraction (Fig. 3, Table 3).

**Figure 3:**
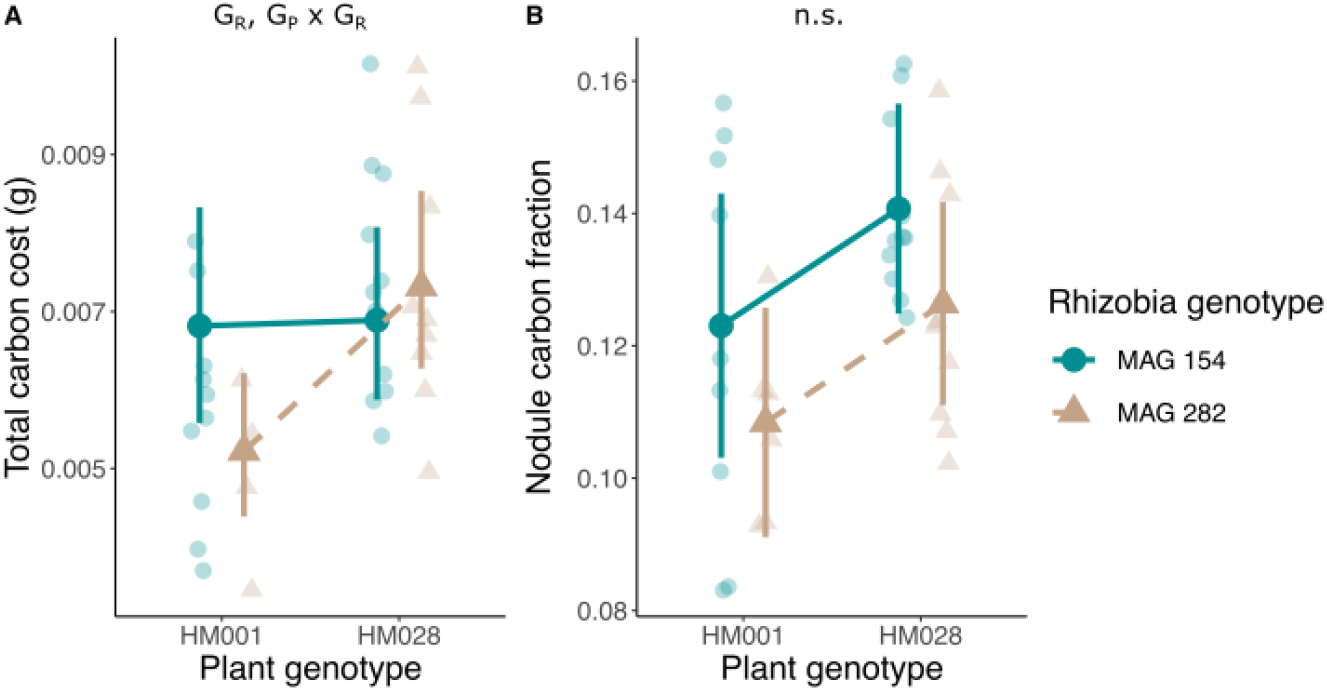
Genetic variation for two cost-associated exchange traits: (A) total carbon cost and (B) the nodule carbon fraction. Transparent points are the raw data and opaque points are the estimated marginal means. Error bars are the 95% confidence interval. Significant genotype effects are listed on the figure: G_P_ = plant genotype effect, G_R_ = rhizobia genotype effect, and G_P_ X G_R_ = plant genotype X rhizobia genotype interaction.

We also quantified the nitrogen-for-carbon exchange ratio by dividing total nitrogen benefit by the total carbon cost (Table 1). We observed a significant rhizobia genotype effect for the nitrogen-for-carbon exchange (Fig. 4, table 3).

**Figure 4:**
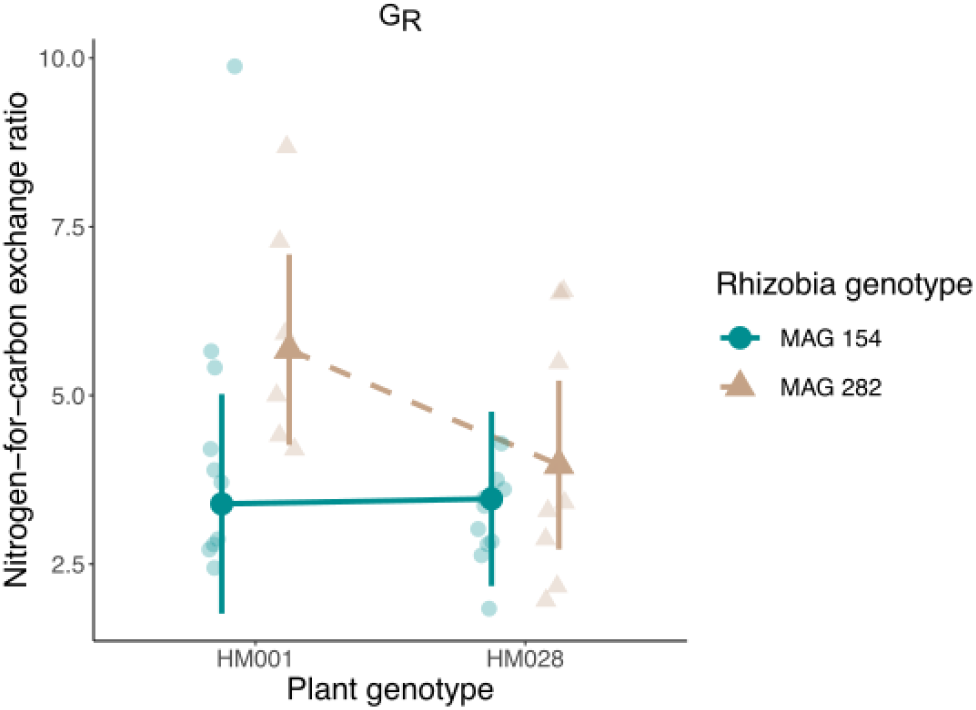
Genetic variation for the nitrogen-for-carbon exchange ratio. **Transparent points are the raw data and** opaque points are the estimated marginal means. Error bars are the 95% confidence interval. Significant genotype effects are listed on the figure: G_**P**_ = planet genotype effect, G_**R**_ = rhizobia genotype effect, and G_**P**_ X G_**R**_ = plant genotype X rhizobia genotype interaction.

### Genetic variation in resource use and production traits

We measured variation in two traits associated with resource use and production, the carbon:nitrogen ratio in aboveground tissue (a proxy for nitrogen use efficiency) and leaf photosynthetic rate (Table 1). The carbon:nitrogen ratio in aboveground tissue was used to infer how efficiently host plants are able to use nitrogen received through trade to incorporate carbon in aboveground tissue (Dovrat et al. 2020). Leaf photosynthetic rate to infer the efficiency of the production of carbon, some of which is ultimately traded away. We found a significant effect of rhizobia genotype and the interaction between rhizobia genotype and plant genotype on carbon:nitrogen ratio (Fig. 5, Table 3). We observed a significant plant genotype x rhizobia genotype interaction on photosynthesis rate (Fig. 5, Table 3).

**Figure 5:**
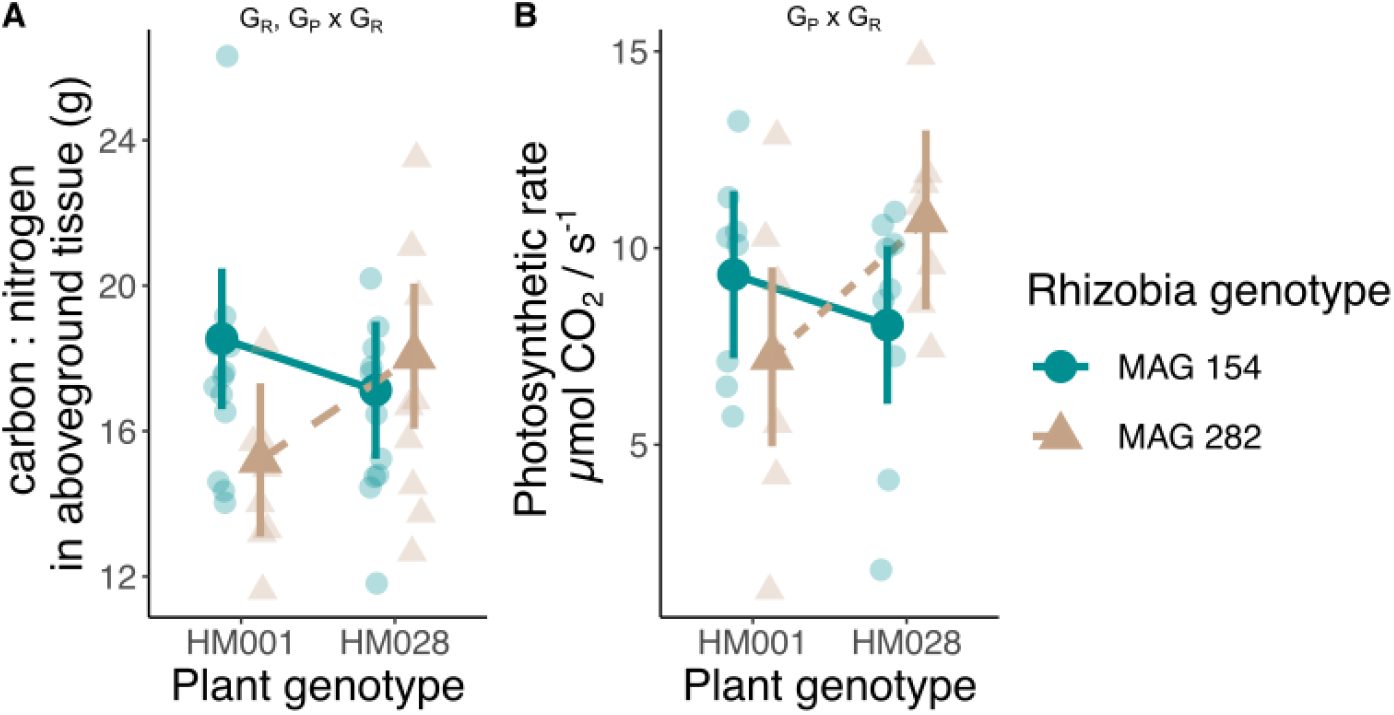
Genetic variation for (A) carbon : nitrogen ratio in aboveground tissue and (B) leaf photosynthetic rate. Transparent points are the raw data and opaque points are the estimated marginal means. Error bars are the 95% confidence interval. Significant genotype effects are listed on the figure: G_P_ = planet genotype effect, G_R_ = rhizobia genotype effect, and G_P_ X G_R_ = plant genotype X rhizobia genotype interaction.

## Discussion

In this study, we found that mutualism resource exchange, use, and production traits are influenced by both plant and rhizobia genotype. Consistent with our expectations, we observed rhizobia genotype and GxG effects on both benefit-associated and cost-associated resource exchange traits. However, we only observed a rhizobia genotype effect for the nitrogen-for-carbon exchange ratio. Contrary to our expectation, rhizobia genotype affected plant resource use and production traits as much or more than plant genotype. However, this result could also be a result of the small number of genotypes we used in our study. Our results suggest that rhizobia genetic variation shapes resource use and production traits in their hosts. Furthermore, our work indicates that the evolution of mutualisms may not be limited to resource exchange traits. Rather, how efficiently mutualistic partners can integrate the benefits of trade (i.e. resource use efficiency) and/or how efficiently they can produce the resource traded away may be just as important for determining the evolution of mutualisms. Together, these results suggest an important role for jointly determined traits in mediating mutualism evolution.

### Genotype effects on resource exchange traits

We predicted that GxG effects, which occur when the genotype effect on trait expression in one partner depends on the genotype of the other partner, would be common to exchange traits because resource exchange involves cooperation between the plant and rhizobia (Sachs et al. 2004). Consistent with this prediction, we observed GxG effects for two benefit-associated traits: nitrogen composition and total nitrogen benefit per symbiont. For cost-associated traits we found GxG effects for total carbon cost. However, we did not observe a GxG effect for the carbon-for-nitrogen exchange ratio. Our prediction that GxG effects should be common to exchange traits is based on the hypothesis that the outcomes of trade in resource mutualisms are governed by bargaining between partners (Sachs et al. 2004; Akçay and Roughgarden 2007). The GxG effects on exchange traits that we observed here could be mediated by the numerous bargaining strategies that hosts and microbes employ. For example, legumes are able to “sanction” poor quality partners resulting in beneficial trading outcomes (Kiers et al. 2003; Westhoek et al. 2021). Alternatively, GxG effects on exchange traits may also emerge via a mismatch between partners (Barrett et al. 2012; Wendlandt et al. 2019). Partner mismatches occur when some rhizobia are more effective partners for some plant genotypes compared to others.

To our surprise, however, the carbon-for-nitrogen exchange rate was only affected by rhizobia genotype, not by both partners. This result suggests that rhizobia may exert more control over the overall outcome of trade than their host. This finding conflicts with theoretical and empirical work that suggests that, in general, host plants exert more power in determining the outcomes of trade (Daubech et al. 2017; Clark et al. 2019). Of course, a primary reason that host plants are expected to exert more control over the trade is their ability to “bargain” with multiple different microbial strains simultaneously (Akçay and Simms 2011; Westhoek et al. 2021); a scenario that was unable to occur in this study due to our single strain inoculation design. It is also possible that the two plant genotypes we used in this experiment do not measurably differ in their ability to manipulate the carbon-for-nitrogen exchange rate and that a larger panel of plant genotypes would have revealed plant genotype variation for resource exchange. Future research must include more plant and rhizobia genotypes to better test the relative control that host plants rhizobia exert over the resource exchange.

Finally, there were a few exchange traits for which we observed no genotype effects. Among benefit-associated traits, total nitrogen benefit, our metric of absolute benefit, did not differ between plant or rhizobia genotypes. The same was true for nodule carbon fraction, our metric of relative cost. The fact that we observed genotype differences in the relative but not absolute benefit—and the absolute but not relative cost—suggests that absolute and relative costs (and benefits) of mutualism may exhibit different evolutionary dynamics. Another explanation is that there is genetic variation for these traits in nature, but was simply not captured by our panel of two plant genotypes and two rhizobia genotypes.

### Genotype effects on resource use and production traits

We predicted that plant genotype would be the sole contributor to differences in plant resource use and production traits. Contrary to this prediction, we observed GxG effects for both of the resource use and production traits that we measured: nitrogen use efficiency (our metric of resource use) and leaf photosynthetic rate (our metric of resource production).

We predicted that host resource use and production traits would respond primarily to plant genotype because these traits are indirectly related to resource exchange and represent components of plant physiology. However, rhizobia effects on these host traits are perhaps not surprising if hosts change how they allocate benefits or invest in the production of resources in response to trade. Thus, rhizobia and GxG effects on resource use and production traits may arise from the GxG effects for exchange traits that we observed. As mentioned above, it is also possible that a lack of plant genotype effect is simply due to a lack of measurable differences in resource use and production traits between the two plant genotypes we used here.

Resource use traits, like nitrogen use efficiency, have previously been shown to be microbially mediated (Dovrat et al. 2020; Zhang et al. 2020). For instance, legumes modify their nitrogen use efficiency in response to both soil nitrogen availability and the presence of rhizobia (Dovrat et al. 2020). Our study raises the important question of why nitrogen use efficiency could vary across combinations of plant and rhizobia genotypes. A GxG effect on nitrogen use efficiency could be due to differences in the efficiency of symbiotic nitrogen fixation across plant and rhizobia combinations. In contrast to some legume species that down-or upregulate nitrogen fixation in response to environmental conditions *M. truncatula* is an obligate fixer that attempts to maintain a constant nitrogen fixation effort regardless of the activity of their rhizobia partners or external supplies of nitrogen (Hedin et al. 2009). Variation in nitrogen use efficiency in obligate fixers like *M. truncatula* could simply be a function of the variation in the efficiency of nitrogen fixation across plant and rhizobia genotype combinations. Alternatively, it is also plausible that legumes may have evolved strategies to adjust how they use and allocate the benefits from trade in order to buffer against the consequences of interacting with poor quality partners.

Resource production traits, like leaf photosynthetic rate, also have been shown to respond to microbes (Adams et al. 2016). Legumes in particular exhibit significantly higher photosynthetic rates in the presence versus the absence of rhizobia (Clark et al. 2019; Dovrat et al. 2020). Photosynthesis is fundamental to the exchange of resources between legumes and rhizobia because it provides the carbon necessary to support rhizobia and it also requires a large amount of nitrogen, which is provided by the rhizobia. A heightened photosynthetic rate may equate to a larger pool of carbon that can be devoted to the mutualism, and by extension, a large supply of nitrogen that can be returned to the host plant. Variation in photosynthetic rate may also simply be an indication of nitrogen limitation and the outcomes of trade.

Photosynthesis involves nitrogen-rich machinery and therefore, genotype combinations with the highest photosynthetic rates may simply be the genotype combinations in which the largest amount of nitrogen is being transferred to the host plant. Regardless of the mechanisms responsible for GxG effects on photosynthetic rate, the fact that a portion of the genetic variation for leaf photosynthetic rate resides in rhizobia emphasizes the far-reaching effect that this mutualism has on host plant physiology.

The findings of this study underscore the possibility that the evolution of mutualisms may involve traits related to how interacting partners can integrate the benefits of trade (i.e. resource use efficiency) and how efficiently they can produce the resources traded away. This suggestion is corroborated by a recent study involving a synthetic mutualism between engineered strains of brewer’s yeast (Vidal and Segraves 2021). The authors observed strong fluctuations in both resource use efficiency and resource production of both strains as they evolved in response to the availability of the limiting resources produced by their partners. Resource use and production traits may represent key components of mutualisms that allow interacting partners the ability to adjust their physiology in response to the outcomes of trade. Future studies should carefully consider selection on all mutualism traits, not just those related to the exchange of resources, as well as covariation among resource exchange, use and production traits.

Resource exchange, use, and production are all dynamic and interrelated processes. An important challenge for the study of mutualisms is to better understand how each of these processes influences the other. The widespread pairwise correlations among the traits we quantified in the study suggests how tightly interwoven these processes are (Fig. S1). They also raise the intriguing possibility that genotype effects on one trait can cause correlated changes in the expression of genetic variation for other traits. However, as stated in the methods, many of the traits we measured here are not mathematically independent owing to the fact that there is some overlap in the variables used to calculate the traits we examined in the study. Future work is needed to carefully isolate the different components of mutualisms to quantify their effects on one another.

### The importance of jointly determined traits to predicting evolution in mutualism

The ubiquity of rhizobia and GxG effects on mutualism traits has three important implications for the evolution of mutualisms. First, this finding indicates that traits involved in resource exchange, use, and production may be jointly determined by the genomes of both interacting partners, just as is the case with fitness outcomes (Heath 2010). Jointly determined traits, here defined as traits that exhibit variation in response to more than one genome as either two independent main effects or a GxG effect, can have different evolutionary trajectories than those determined by the genome of just one or the other interacting partner (O’Brien et al. 2021b). Thus, a trait-based framework, wherein quantitative genetic approaches can be leveraged to identify the sources of variation in mutualism traits, is necessary to develop a better understanding of mutualism evolution.

Second, these results add to the growing body of literature demonstrating that microbial mutualists play an important role in the expression of host traits like resource use and production. Microbes are known to mediate host nutrient acquisition (Hestrin et al. 2019; Harbort et al. 2020), life history transitions (Wagner et al. 2014; O’Brien et al. 2021a), defense against antagonists (Johnson et al. 1987; Berg and Koskella 2018), stress tolerance (Lau and Lennon 2012), and morphology (Ortíz-Castro et al. 2008; Batstone et al. 2017). Our work builds on this knowledge by demonstrating that intraspecific variation in microbial populations, in addition to interspecific variation, can also influence the expression of plant functional traits. This finding complements two recent studies, O’Brien et al. (2021b) and De Lisle et al. (2022), which have highlighted the potential evolutionary significance of the effect of intraspecific partner variation in the expression of mutualism traits. Crucially, intraspecific microbial variation is invisible to the 16S sequencing that is commonly applied to characterize community composition in microbiome studies because strains belonging to the same species have identical 16S sequences. Thus, substantial cryptic variation in microbe-mediated plant traits measured in microbiome manipulations may go unnoticed.

Third, variation in resource trade, use, and production traits across different combinations of plant and rhizobia genotypes highlights the potential importance of the non-random association of partner genotypes for mutualism evolution. If a mutualism trait responds to a GxG effect and is a target of selection, then the plant genotypes favored by selection will be dependent on which rhizobia genotype they are paired with. If associations between specific plant and rhizobia genotypes are random, then the response to selection will be smaller than expected because most genetic variation is non-additive and a function of which genotypes are paired together. On the other hand, if associations between specific plant and rhizobia genotypes are non-random, then the response to selection could be either larger (if associations between plant and rhizobia genotypes inflate genetic variation for the trait) or smaller (if associations between plant and rhizobia genotypes constrain genetic variation for the trait) than expected. While host plants have a clear influence on their rhizosphere microbes (Miranda-Sánchez et al. 2016; Greenlon et al. 2019) and can vary in their ability to select beneficial microbial genotypes (Burghardt et al. 2018, 2019), the frequency of non-random associations between specific plant and rhizobia genotypes in nature has been rarely measured. Given the apparent importance of rhizobia genotypes in influencing variation in mutualism traits, more studies are needed to address the degrees of non-random associations between interacting genotypes in order to develop a predictive framework for understanding mutualism evolution.

Ultimately, the relative importance of genotype and GxG variation for mutualism traits to mutualism evolution is dependent on the contribution of that variation to fitness. An important next step to expand our understanding of mutualisms is to quantify the relationship between resource exchange, use, and production traits and fitness, i.e. selection. While we did measure common fitness proxies for this system, such as plant aboveground biomass, these fitness proxies were also used in the calculation of most of the resource exchange, use, and production traits presented in this study. The mathematical non-independence between our fitness proxies and resource exchange, use, and production traits precluded us from being able to measure selection. Measurements of selection will reveal both the traits that underpin mutualism variation and will allow us to begin dissecting the selective agents that maintain fitness variation in mutualisms.

## Acknowledgements

We thank Chang-Yu Chang, Addison Martin, and Eunnuri Yi for helpful comments on this manuscript. We also thank Chigozie Ibe, Anisa Robinson, Linda Wu, and Seokyoon Chang for help with executing the experiment and collecting data. *Medicago truncatula* genotypes were generously provided by Nevin Young and the Medicago HapMap project. *Ensifer meliloti* strains were generously provided by Liana Burghardt and Katy Heath. Growth chamber logistical support was provided by the University of Pennsylvania’s Greenhouse support staff members Kathyrn Butler and Samara Gray. We are grateful to David Vann and the University of Pennsylvania’s Earth and Environmental Science Department’s Instrumentation Core for assistance with elemental analysis. This work was funded by a Society for the Study of Evolution R. C. Lewontin Award to McCall Calvert and NSF-DEB 2118397 to Corlett Wood

## Notes

### Competing Interest Statement

The authors have declared no competing interest.

### Summary of Updates

The scope and research objectives of the manuscript were revised to better align with the results that are presented.

