## Supplemental information for "Genotypic variation in resource exchange, use, and production traits in the legume-rhizobia mutualism"

**Supplementary Information**

**Methods**

*Photosynthesis rate measurement and estimation*

We performed photosynthesis measurements on two or three plants from each genotype combination at 10 weeks post planting. Measurements were made using a 6400-40 LCF leaf chamber and 6400-02B LED Light Source connected to a Li-Cor LI-6400 apparatus (LI-COR, Inc., Lincoln, NE, USA). The leaf chamber was illuminated with LED lights emitting at at PAR of ~2000 μmol/$m^{-2}$/$s^{-1}$ and provided with air containing 400 μmol $CO_2$ per mol of air at a constant airflow of 500 μmol $s^{-1}$. The Li-Cor LI6400 estimates photosynthesis by measuring differences in the the amount of $CO_2$ entering and leaving the chamber using infrared gas analysis. The photosynthesis rate was allowed to equilibrate for roughly 10 to 15 minutes prior to each measurement. We measured three leaves on each plant.

We estimated photosynthesis rate as the μmol $CO_2$ $m^{-2}$ $s^{-1}$. Leave level μmol $CO_2$ $m^{-2}$ $s^{-1}$ was estimated using the equation $A = \frac{F\left(C_r - C_s\left(\frac{1000-W_r}{1000-W_s}\right)\right)}{100S}$, where A is the assimilation rate (μmol $CO_2$ $m^{-2}$ $s^{-1}$), F is the molar flower rate of air entering the leaf chamber (this was set to 500 μmol $s^{-1}$), $C_r$ is the the mole fraction of $CO_2$, μmol $CO_2$ $mol^{-1}$ air, measured with the reference infrared gas analyzer (IRGA)(this value is calibrated to 400 ppm), $C_s$ is the mole fraction of $CO_2$, μmol $CO_2$ $mol^{-1}$ air, in the sample IRGA, $W_r$ is the is the mole fraction of water vapor, mmol $H_2O$ mol $air^{-1}$, in the reference IRGA, $W_s$ in the mole fraction of water vapor, mmol $H_2O$ mol $air^{-1}$, in the sample IRGA, and S is the leaf area, $cm^2$. Leaf area estimated by calculating the surface area of scanned images of leaves in imageJ.

*Seed germination*

We broke open seed pods to free seeds by placing seed pods in between two pieces of corrugated plastic and rubbing the pieces together. The seeds were then manually scarified with sandpaper, sterilized with 10% bleach, and placed on moistened filter paper in Petri dishes sealed with Parafilm™ (Bemis Company, Neenah, WI, USA). We covered the sealed Petri dishes with aluminum foil to prevent light from inducing chlorophyll activation and moved them to a 4 C refrigerator. After two days at 4 C we removed the plates from the fridge, inverted them to promote radicle elongation, and placed them in a cabinet at room temperature. After two days, we removed the aluminum foil and placed the petri dishes on the bench top. Once the cotyledons began to turn green (approximately three hours) we started planting.

### Supplementary Figures

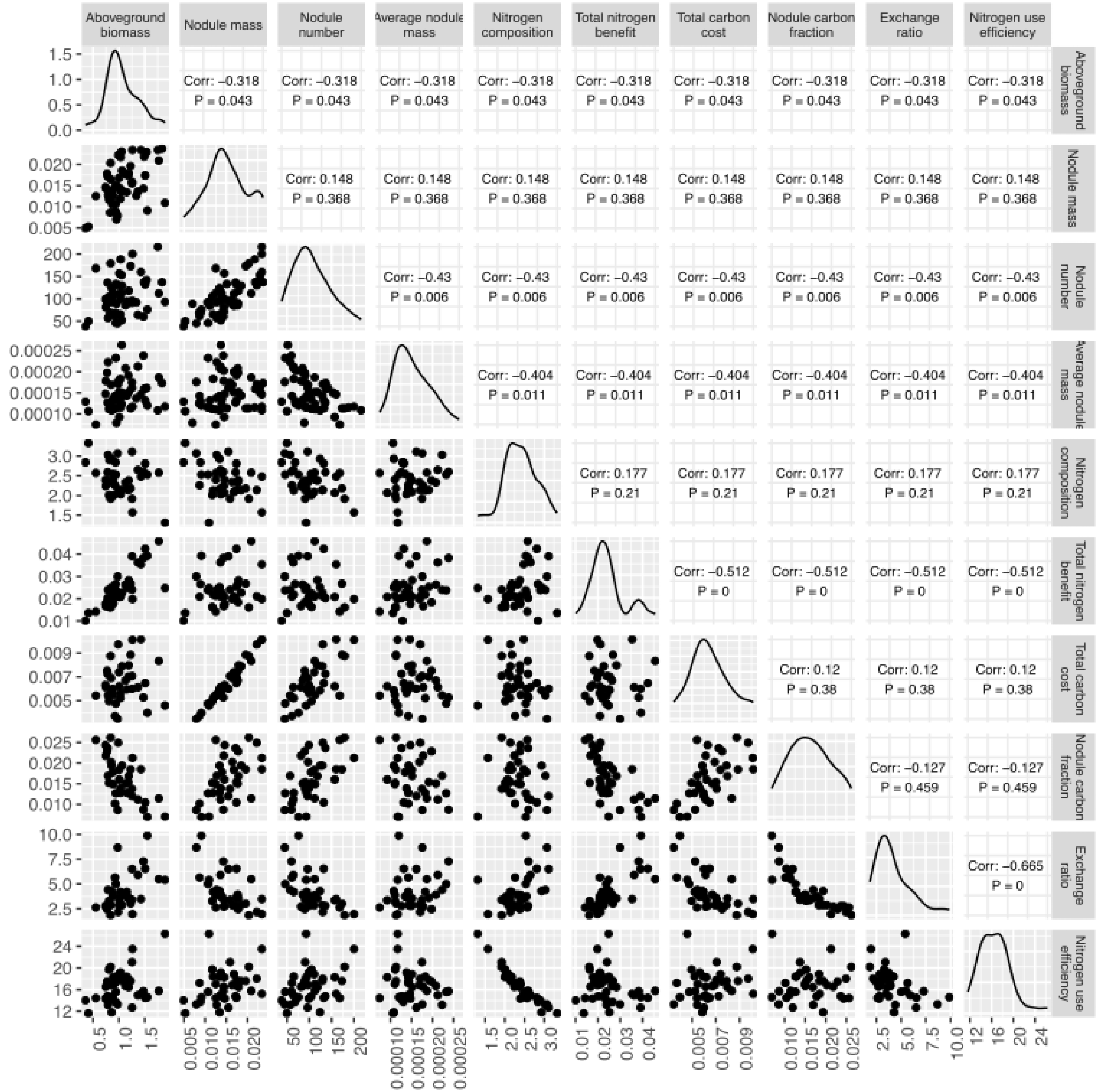


**Supplementary figure 1.** Correlogram of resource exchange and use traits. The bottom diagonal portion of the correlogram consists of plots of the raw values of the two traits being compared. The upper diagonal portion consists of the Pearson correlation coefficient and Bonferroni adjusted p-values for each trait correlation test.
